# Conserved function of ether lipids and sphingolipids in the early secretory pathway

**DOI:** 10.1101/2019.12.19.881094

**Authors:** Noemi Jiménez-Rojo, Manuel D. Leonetti, Valeria Zoni, Adai Colom, Suihan Feng, Namrata R. Iyengar, Stefan Matile, Aurélien Roux, Stefano Vanni, Jonathan S. Weissman, Howard Riezman

**Affiliations:** NCCR Chemical Biology, Department of Biochemistry, University of Geneva, 1211, Geneva, Switzerland; Biofisika Institute (CSIC, UPV/EHU) and Department of Biochemistry, University of the Basque Country, Leioa, Spain; Department of Cellular and Molecular Pharmacology, University of California, San Francisco, California 94158; Howard Hughes Medical Institute, San Francisco, California 94158; Department of Biology, University of Fribourg, Fribourg, Switzerland; Institute of Protein Biochemistry (IBP), Italian National Research Council (CNR), Napoli, Italy; NCCR Chemical Biology, Department of Organic Chemistry, University of Geneva, 1211, Geneva, Switzerland

## Abstract

Sphingolipids have been shown to play important roles in physiology and cell biology, but a systematic examination of their functions is lacking. We performed a genome-wide CRISPRi screen in sphingolipid-depleted cells and identified hypersensitive mutants in genes of membrane trafficking and lipid biosynthesis, including ether lipid synthesis. Systematic lipidomic analysis showed a coordinate regulation of ether lipids with sphingolipids, where depletion of one of these lipid types resulted in increases in the other, suggesting an adaptation and functional compensation. Biophysical experiments on model membranes show common properties of these structurally diverse lipids that also share a known function as GPI anchors in different kingdoms of life. Molecular dynamics simulations show a selective enrichment of ether phosphatidylcholine around p24 proteins, which are receptors for the export of GPI-anchored proteins and have been shown to bind a specific sphingomyelin species. Our results support a model of convergent evolution of proteins and lipids, based on their physico-chemical properties, to regulate GPI-anchored protein transport and maintain homeostasis in the early secretory pathway.

## INTRODUCTION

The maintenance of membrane lipid homeostasis is an energetically expensive yet necessary process in cells. Lipid diversity has evolved together with cell complexity to give rise to thousands of different lipids species with specific functions, many of which are still unexplored ^1, 2^. Moreover, different lipid metabolic pathways are interconnected, and cells show a high phenotypic plasticity when adapting to changes in membrane lipid composition, which makes it difficult to disentangle the function of individual lipid species ^3^. A systematic analysis of the cellular responses to perturbation of specific synthetic pathways is thus needed to reveal co-regulated lipid networks and uncover new lipid functions.

Sphingolipids (SL) are a class of lipids that contain a sphingoid-base backbone, in contrast to the more commonly found glycerol backbone in glycerophospholipids (GPL). These bioactive lipids have been extensively studied in the last decades, revealing distinctive physico-chemical properties and connections to diseases ^4, 5^. SL have been implicated in diabetes ^6^, cancer ^7^ and inflammation ^8^, and mutations in SL synthetic or metabolic enzymes are associated with severe genetic disorders ^9-11^. SL species sphingosine (So) and ceramide (Cer) can permeabilize membranes ^12, 13^; Cer induces the flip-flop of neighbouring lipids ^14^ and can phase-separate to form membrane platforms important for signalling ^15^. The most abundant SL species, sphingomyelin (SM), has been shown to modulate membrane properties and regulate signalling pathways ^16, 17^. Besides direct phosphorylation of ceramide synthases ^18-20^, the only direct regulators of sphingolipid synthesis identified are Orm proteins (ORMDL in mammalian cells), that associate with serine palmitoyl transferase (SPT), the first enzyme of the sphingolipid *de novo* synthetic pathway, and mediate the feedback regulation of sphingolipid production in cells ^21, 22^. Despite this body of work, many basic aspects of the regulation, molecular interactors and functions of SL still remain poorly understood.

Ether lipids (EL) form a poorly characterized lipid family which represents a subclass of glycerophospholipids that are defined by the presence of an ether bond (or vinyl-ether, for plasmalogens) in lieu of the ester bond found in canonical phospholipids at the sn-1 position. This feature, which is intermittently represented across the evolutionary tree, is decisive for the behaviour of these lipids in membranes, and for specific lipid-protein interactions ^23^. Interestingly, EL are not only present in membranes as abundant lipid components, where they can range from approximately 5 to 20 mol % of the total GPL content, but are also a major building block of glycosylphosphatidylinositol (GPI) anchors, a specialized protein post-translational modification. Peroxisomal synthesis of EL is necessary for the synthesis of the 1-alkyl, 2-acyl phosphatidylinositol GPI anchor, which is attached to most of the mammalian GPI-anchored proteins (GPI-AP) in the ER ^24, 25^. GPI-AP are then exported (through binding to p24 proteins: TMED2 and TMED10 in human) from the ER to the plasma membrane via the Golgi complex, where they undergo different remodelling modifications replacing the unsaturated fatty acid in the sn-2 position with a saturated one. Interestingly, the same p24 proteins that selectively pack GPI-AP into the COPII carriers ^26, 27^ are also involved in COPI-dependent retrograde transport ^28, 29^ and p24 proteins bind a particular sphingomyelin species (SM C18 n-acyl) that regulates the initiation of COPI vesicle budding and retrograde transport from the Golgi to the ER^30^.

Here, we apply high-throughput genetics and lipidomic methods to uncover novel regulatory aspects of sphingolipid metabolism and we find a co-regulation of this lipid class with the structurally unrelated EL. Our multidisciplinary approach combining MD simulations with the study of GPI-AP transport kinetics establishes an active role for (ether)lipids in the transport of GPI-AP from the ER to the Golgi, and emphasizes the physico-chemical properties of EL that enabled them to be selected in mammalian cells to perform this function.

## RESULTS AND DISCUSSION

To identify pathways regulating cellular responses to sphingolipid depletion, we performed a genome-wide CRISPRi growth screen for genes that modulate sensitivity to myriocin, an inhibitor of *de novo* sphingolipid synthesis. To insure dependence on *de novo* lipid synthesis (as opposed to exogenous lipid intake, which can circumvent the need for cellular synthesis ^31^), we cultured human K562 cells in lipid-depleted serum (see Material and Methods). Indeed, we found that myriocin treatment efficiently arrested growth in lipid-depleted media but not in lipid-rich media, while lipid depletion alone did not compromise growth (Fig S1A). Lipidomic analysis verified that myriocin treatment severely decreased cellular sphingolipid levels (Fig 2 A, B, Fig S1 B) under these conditions.

**Figure 1.**
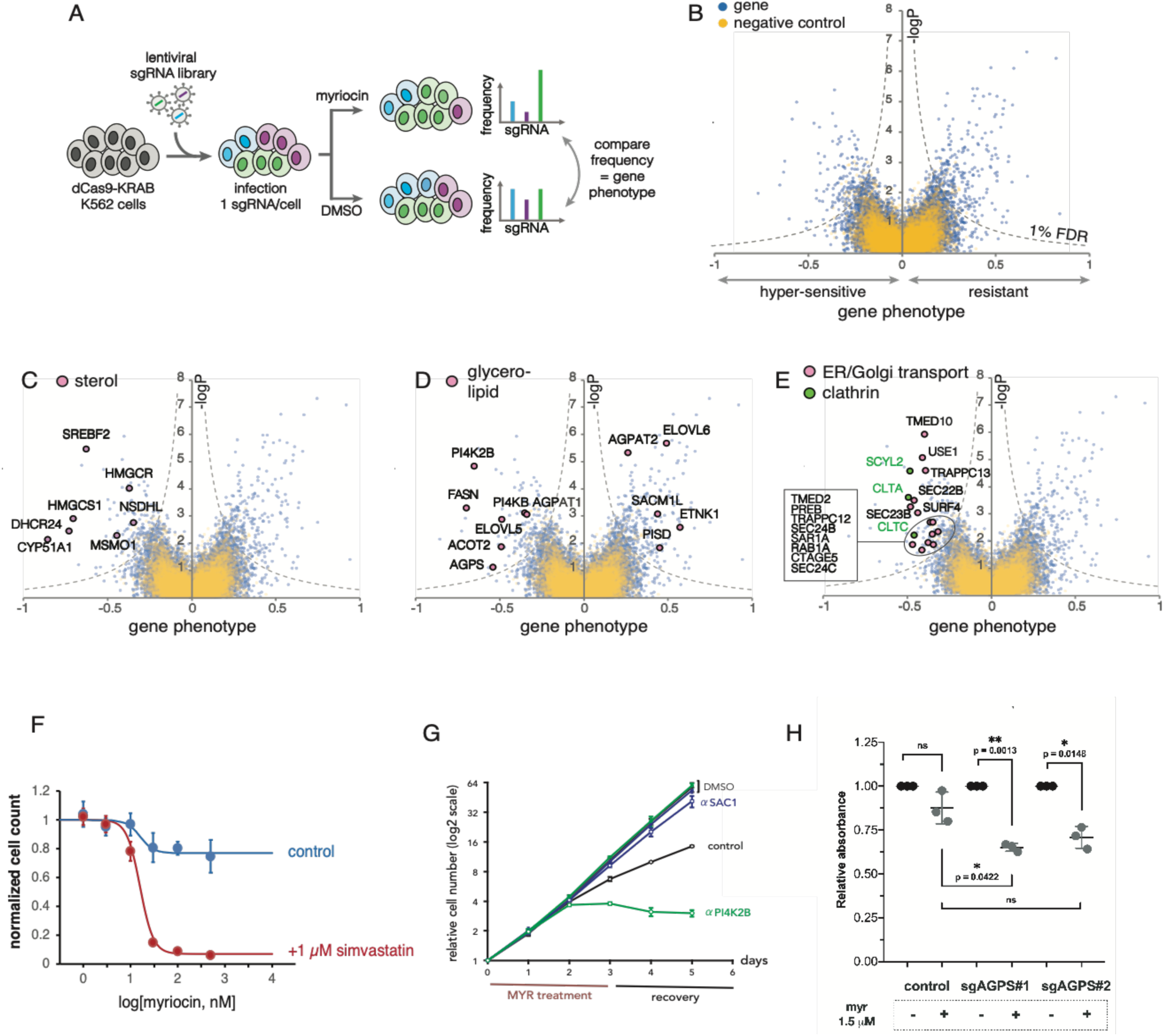
A genetic screen in sphingolipid-depleted cells highlights lipid co-dependencies and uncovers a major role for sphingolipids in intracellular transport. **a**, Schematic of myriocin sensitivity CRISPRi screen in K562 cells. **b**, Volcano plot showing gene phenotype on the X-axis and logP value on the Y-axis. Each dot is an individual gene (blue). Negative control phenotypes reconstructed from non-targeting sgRNAs are shown in yellow. **c**, Hits that include genes involved in sterol metabolism. **d**, Hits involved in membrane lipid metabolism. **e**, Hits involved in intracellular transport including those involved in the early secretory pathway and clathrin mediated trafficking. **f**, Myriocin dose-response curves showing K562 cell survival in the absence (control) or presence of 1 *µ*M simvastatin after 3 days of treatment in de-lipidated media. Each point represents average ± standard deviation from 3 independent measurements (some error bars are smaller than symbols). Sigmoid fits (solid lines) with IC50 = 16 nM are shown. **g**, Myriocin sensitivity of CRISPRi K562 dCas9-Krab lines. Growth curves are shown for parental line (no sgRNA, “control”), SACM1L knock-down (aSAC1) and PI4K2B knock-down (aPI4K2B). Note that cell count (y-axis) is shown on a log2 scale. Open symbols: growth upon 3-day treatment with 1*µ*M myriocin, followed by 3 days of recovery. Full symbols: un-treated controls (DMSO). SACM1L knock-down protects from the growth impact of myriocin, while PI4K2B knock-down accentuates its effect. Shown are average and standard deviation from 3 independent replicate experiments. **h**, Myriocin sensitivity of ether lipid-depleted Hela CRISPRi cell lines using the colorimetric MTT assay. Data were normalized to control for each condition (vehicle). Each data point represents the average absorbance in one well.

**Figure 2.**
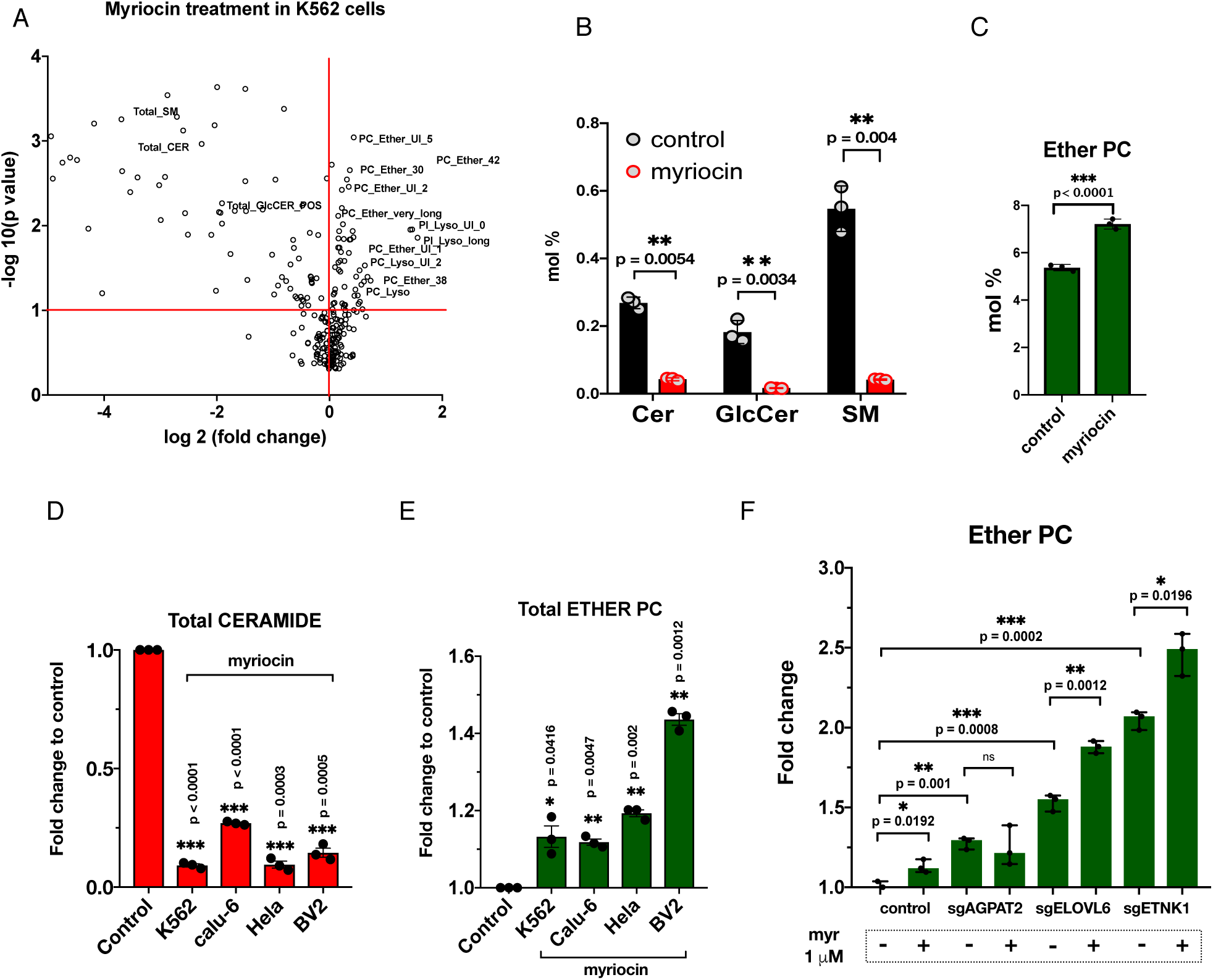
Systematic lipidomic analysis reveals a metabolic co-regulation of sphingolipids and ether lipids. **a**, Lipid changes in K562 cells after 3-day inhibition of sphingolipid synthesis with 1 *µ*M myriocin. Log 2 of the fold change compared to control cells in the X-axis and the -log P value in the Y-axis (n=3). **b**, Relative sphingolipid species amount over the total of lipids detected (n=3). Cer: total ceramide; GlcCer: total glucosylceramide; SM: total sphingomyelin. **c**, Relative ether phosphatidylcholine (PC) levels (n=3). **d**, Decrease in ceramide levels upon myriocin treatment in different cell lines (n=3). **e**, Increase in ether PC levels upon myriocin treatment in different cell lines (n=3). **f**, Relative ether PC levels in myriocin-resistant cell lines before and after treatment (n=3). Each condition represents the mean of 3 independent measurements of approximately 2 million cells. Statistical analysis was performed by unpaired two-tailed Student’s t-test.

To systematically repress gene expression, we used an ultra-complex genome-wide CRISPRi library targeting 15,977 genes using 10 sgRNAs per gene ^32^, also including 1,000 non-targeting control sgRNAs to allow precise measurement of experimental noise. For screening, K562 cells stably expressing dCas9-KRAB were infected with a pooled lentiviral library of sgRNA constructs, and dCas9/sgRNA expressing cells were treated with myriocin or DMSO in lipid-depleted media (Fig 1 A). Deep sequencing analysis using sgRNA sequences as barcodes ^32, 33^ revealed the impact of individual gene knock-down on survival under sphingolipid restriction. Screening analysis revealed a rich dataset, with over 450 genes regulating survival under myriocin treatment (Fig 1 B). Strikingly, several gene families emerged with strong phenotypes, linking survival under sphingolipid depletion to other lipid synthetic pathways and intracellular vesicular transport in particular.

First, repression of sterol synthesis enzymes or their transcriptional activator SREBF2 strongly compromised the ability of cells to survive without SL (Fig 1 C). We further verified this strong co-dependency between SL and sterols by showing that sterol synthesis inhibition with simvastatin greatly accentuates the impact of myriocin on growth (Fig 1 F). This highlights the synergistic function of sphingolipid and sterols which has been previously described in yeast by Guan *et al*., where the authors found correlative changes in sterol and sphingolipid homeostasis and synthetic phenotypes between different sterol and sphingolipid mutants ^34^, demonstrating that sterols and sphingolipid work together in biological membranes.

Second, perturbations of different routes of glycerophospholipid metabolism and remodeling showed both aggravating or resistant phenotypes (Fig 1 D), indicating a functional cross-talk between SL and specific GPL species. Interestingly, specific pathways for acyl chain remodeling showed either resistance or hyper-sensitivity to SL depletion (ELOVL5/6, AGPAT1/2). In particular, repression of ELOVL5, an enzyme with specificity to elongate polyunsaturated fatty acids (PUFA), is hypersensitive, while ELOVL6, which has specificity to elongate saturated C16 acyl CoAs is a resistant hit, showing that acyl chain saturation can modulate survival in the absence of SL synthesis, suggesting that an increase in the PUFA to saturated fatty acids ratio aids growth under sphingolipid depletion. Our results also highlighted a strong interaction between SL and phosphatidylinositol-4-phosphate (PI4P) synthesis. Indeed, repression of PI4-kinases (PI4K, PI4K2B) strongly compromised growth while repression of the PI4P-phosphatase SACM1L (SAC1) allowed survival upon sphingolipid depletion. This striking signature was verified in individual follow-up experiments (Fig 1 G), and mirrors previous results in *S. cerevisiae* where Sac1 knock-out renders yeast resistant to myriocin (Breslow et al 2010), suggesting that the link between sphingolipid and PI4P homeostasis has been conserved during evolution. Indeed, SL have also been shown to regulate phosphoinositide turnover at the Golgi in mammalian cells, and the metabolic coupling of PI4 kinases/Sac1 phosphatase seems to be responsible for the control of phosphoinositide phosphate (PtdIns(4)P) amounts in response to sphingolipid levels in the Golgi ^35^.

Interestingly, inhibition of ether lipid synthesis through repression of dihydroxyacetone-phosphate synthase (AGPS), a peroxisomal enzyme responsible for the first ether-linked precursor in EL synthesis also compromised survival to myriocin treatment. This functional interaction was confirmed in another cell line: AGPS knock-down in CRISPRi HeLa cells using two sgRNAs also displayed myriocin hyper-sensitivity (Fig 1 H). The first guide RNA (sgAGPS#1) was used for further experiments throughout the paper as it showed the best knock-down efficiency and the strongest decrease in ether lipid levels (Fig S1 D, E). Altogether, our genetic screen revealed rich functional cross-talks between SL and a variety of other lipid species (in particular sterols, PI4P and EL).

Finally, sphingolipid-depleted cells were particularly vulnerable to inhibition of intracellular vesicular transport (Fig 1 E). Specifically, many members of the COPII machinery (responsible for ER to Golgi transport), including coat proteins (PREB (SEC12), SAR1, SEC24B/C, SEC23B), cargo receptors (TMED2, TMED10) and proteins involved in the docking and fusion of the carriers (RAB1A, SEC22B, USE1) were strong hyper-sensitive hits, as were key proteins in clathrin-mediated transport (CLTA, CLTC, SCYL2). SCYL2 has been proposed to regulate clathrin-mediated transport from TGN to endosomes ^36^, suggesting that maintenance of SL levels are specifically important for traffic through the secretory pathway. Some of these hits raise new questions in the sphingolipid and membrane trafficking fields. For instance, cells lacking SURF4, a protein affecting ER morphology are hypersensitive to myriocin, as well as cells knocked-down for cTAGE5, a protein involved in the ER to Golgi transport of large cargos such as collagen ^37^. Elucidating the mechanism by which SL regulate secretion of large cargos could be of interest to understand collagen-related diseases among others ^38, 39^.

Because our screen revealed strong functional interplays between SL levels and other steps in lipid metabolism, we decided to combine the genetic data with lipidomic analysis to identify co-regulated lipid species and new lipid functions. This analysis highlighted a pronounced crosstalk between SL and EL levels. First, when profiling the lipidome of sphingolipid-depleted cells after myriocin treatment, we detected a consistent increase in EL, in particular ether PC species (Fig 2 A, C, Fig S1 B, D). Indeed, we could see this metabolic response in cell lines from different tissues, showing a conserved response including mouse and human cell lines (Fig 2 D, E). Furthermore, lipidomic analysis of some of the resistant hits from the CRISPRi screen (Fig 1 D) showed that they had higher amounts of ether PC in comparison with control cells (Fig 2 F), suggesting that having increased baseline levels of EL can make cells resistant to sphingolipid depletion.

Overall, these results show a co-regulation between the levels of EL and SL in mammalian cells that can explain the genetic interaction previously seen in our myriocin screen. This prompted us to investigate whether the response of cells to increased ether lipid levels upon sphingolipid depletion was bidirectional. For that purpose, we used AGPS knock-down cells and analyzed their lipidome to see if those cells, depleted of EL, showed changes in SL levels (Fig 3 A-G). Indeed, under these conditions the total amount of Cer was increased around 20% driven by increases in a wide range (C16, C18, C22 and C24) of ceramide species (Fig 3 D), while glucosylceramide (GlcCer) levels were unchanged (Fig 3 A). Moreover, even though the total amount of SM and the levels of the major SM (SM C24:1) were not affected, we found a decrease in a specific SM species, SM with a C18 n-acyl chain (Fig 3 D, E).

**Figure 3.**
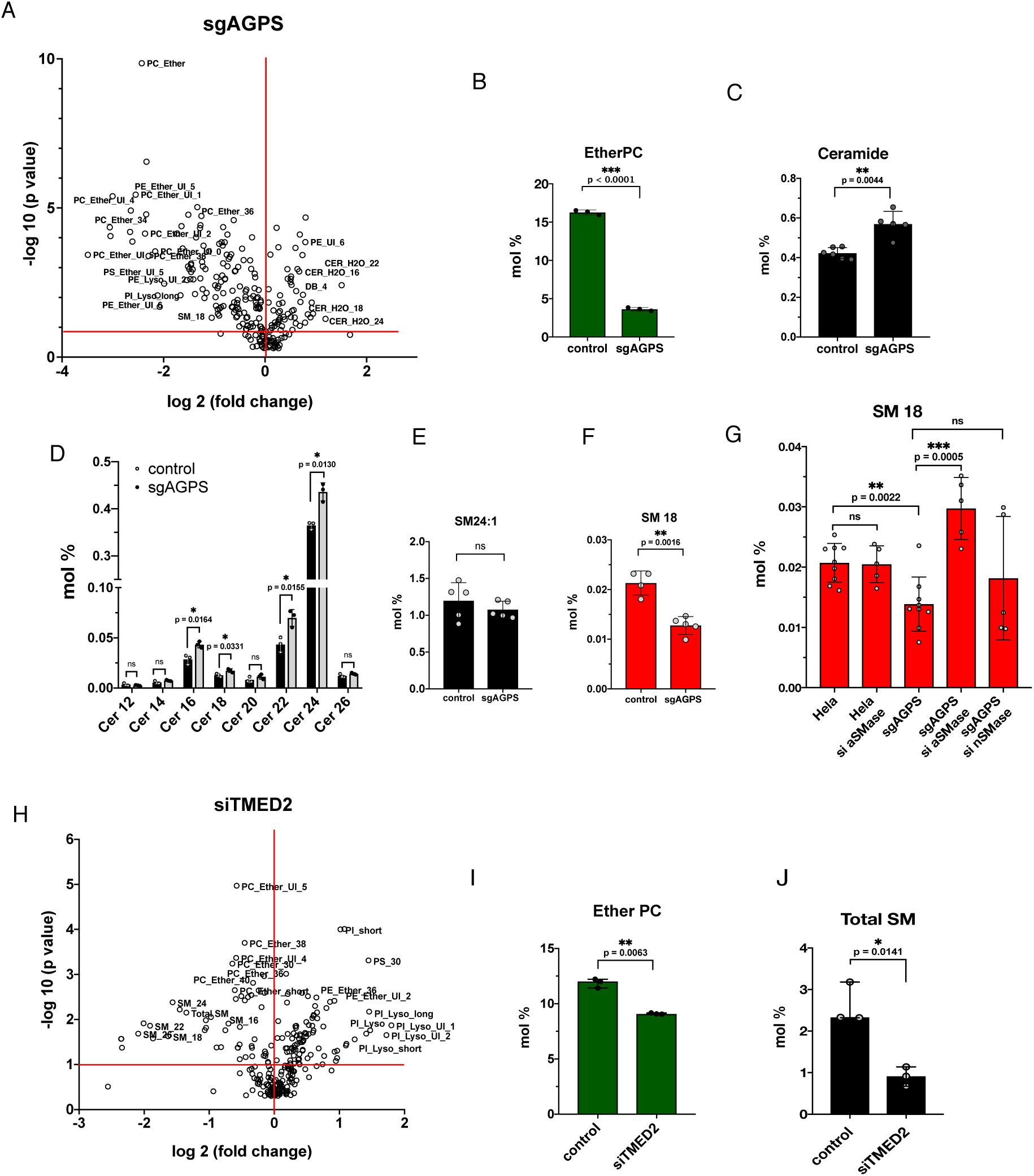
Lipidomic analysis of ether lipid-deficient cells confirms the crosstalk between sphingolipids and ether lipids. **a**, Lipid changes in ether lipid-deficient Hela cells where alkyldihydroxyacetonephosphate synthase (AGPS) has been knocked-down using CRISPRi. Log 2 of the fold change compared to control cells in the X-axis and the -log P value in the Y-axis. **b**, Relative ether PC levels over the total of lipids detected (n=3). **c**, Relative ceramide levels over the total of lipids detected. Cer: total ceramide(n=5). **d**, Relative SM 24:1 levels over the total of lipids detected (n=5). **e**, Relative SM C18 levels over the total of lipids detected (n=5). **f**, Ceramide profile of ether lipid-depleted cells (n=3). **g**, Relative SM C18 levels in ether lipid-deficient cells are rescued by the knockdown of the lysosome-localised acid sphingomyelinase (aSMase, SMPD1) but not by the neutral sphingomyelinase (nSMase, SMPD3). Cells were transfected with siRNA for 3 days and then analysed by tandem mass spectrometry. **h**, Lipid changes in p24 (TMED2) knock-down Hela cells. Log 2 of the fold change compared to control cells in the X-axis and the -log P value in the Y-axis. **i**, Relative ether PC levels in p24 (TMED2) knocked-down cells (n=3). **j**, Relative SM levels in p24 (TMED2) knocked-down cells (n=3).

A unifying interpretation of these lipid changes suggested a possible role of EL in the early secretory pathway. First, this specific SM (SM C18) has been shown to be important for an efficient COPI-dependent retrograde transport from Golgi to the ER through specific binding to p24 proteins ^30^, receptors for the export of GPI-AP ^27^. Interestingly, we find some components of the p24 complex as hypersensitive in the myriocin screen (TMED2, TMED10) (Fig 1 E) and cells lacking TMED2 show a decrease in the levels of EL and SM (Fig 3 H-J), which shows how the levels of these lipids respond to defects in the secretory pathway.

It is also possible that changes in the levels of SM C18 could be used as an indicator of defects in COPI-dependent retrograde transport and that, when retrograde transport is reduced, SM C18 is redirected to the endosomal-lysosomal pathway. To test this, we attempted to rescue the decrease in SM C18 levels in AGPS KD cells by knocking-down the acid sphingomyelinase (aSMase), which is located in the lysosome (Fig 3G). Indeed, reducing aSMase, but not the neutral sphingomyelinase, restored levels of SM C18.

The genetic interaction seen between EL and SL metabolism in the screening, and the bidirectional co-regulation found in our lipidomics data between ceramide and ether PC, indicated that even though these lipids are very different in terms of their chemical structure, they might share functions in cell membranes so that cells could adapt to the lack of one of them by upregulating the other class. Indeed, one common function, apparent from different kingdoms of life, is as part of the lipid anchor in GPI-AP; in yeast the plasma membrane GPI-AP have a ceramide-based anchor ^40^ while in mammalian cells most have an ether lipid-based anchor ^41^. Therefore, we speculated that SL and EL could share functions in the secretory pathway and that their selection for this function could be based on their physico-chemical properties and behavior in membranes

In order to investigate the functional interaction of SL and EL in mammalian cells, and to understand why EL replaced Cer as the lipid moiety of GPI-AP during evolution, we considered the possibility that EL and ceramides might share physico-chemical properties. To investigate this, we used a simplified system and reconstituted bilayers composed of ether PC or Cer in combination with other lipids, in particular phosphatidylcholine (POPC), that we used as a model “scaffold” for the ER membrane (Fig 4 A). To probe lipid packing of these giant unilamellar vesicles (GUVs) we used a mechanosensitive membrane probe (FLIPPER-TR®) recently described to measure membrane tension and lipid packing ^42, 43^. Both ether PC and Cer (at 5 mol % of Cer 24:1 where microscopic phase separation was not detectable) showed similar behavior, decreasing the lifetime of the FLIPPER-TR® probe when present in GUVs composed of POPC (Fig. 4 B). Ether PC alone showed an even more marked decrease in the fluorescence lifetime of the probe. Interestingly, Cer had no effect when present in GUVs composed of ether PC (Fig 4 B). This difference in behavior could be explained by a differential interaction of Cer molecules with ether PC compared to POPC, a hypothesis that is supported by a slight decrease in the amount of hydrogen bonds that ceramide forms with ether PC (Fig S2 A) compared to POPC and a small difference in interlipid distance (Fig S2 B). Thus, even though they are not microscopically visible with this lipid composition, both Cer 24:1 and ether PC could form nanoscopic packing defects in POPC membranes, and the lipid decompression in those areas would be detected as a decrease in fluorescent lifetime of the probe due to a deplanarization of the FLIPPER-TR® molecule ^42^. A similar behavior of Cer and ether PC in the membrane was confirmed by MD simulations, as 2D density maps of bilayers composed of a mixture of POPC: ether PC 85:15 or 70:30 showed a heterogeneous distribution of the two lipids, that is reminiscent of that of POPC-CER mixtures (Fig 4 C).

**Figure 4.**
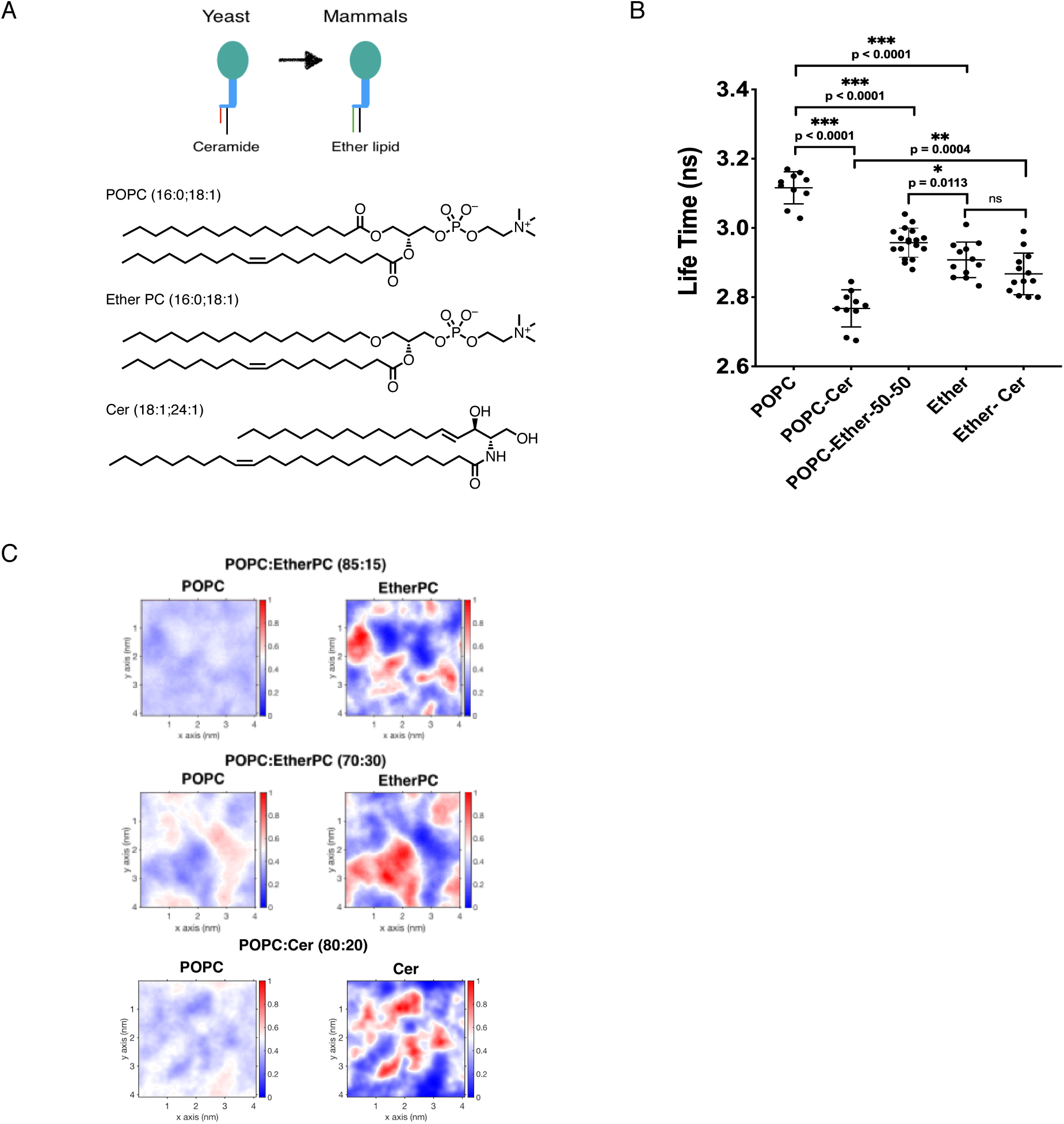
Similar effects of ceramide and ether lipids on membrane organization. **a**, Upper panel: schematic representation of GPI-AP structure in yeast (ceramide based anchor) and in mammalian cells (ether lipid based anchor). Lower panel: structure of the lipids used for this study. POPC, 1-palmitoyl-2-oleoyl-glycero-3-phosphocholine; Cer 24:1, N-nervonoyl-D-erythro-sphingosine; Ether PC, 1-O-hexadecyl-2-oleoyl-sn-glycero-3-phosphocholine. **b**, Fluorescence lifetime values of the FLIPPER-TR® probe as a function of lipid composition in giant unilamellar vesicles. GUVs were composed of either POPC or ether PC and binary mixtures of POPC + 5% ceramide 24:1 and ether PC + 5% ceramide 24:1 as well as an equimolar mixture of POPC and ether PC. **c**, Density maps from MD simulations of the distribution of POPC and either Ether PC or Cer in systems at different concentrations. Red corresponds to enrichment, while blue corresponds to depletion of the lipids.

These results show that ether PC segregates from POPC in a similar way as ceramide, and that membranes composed of ether PC are less packed than pure POPC bilayers. In summary, ether PC and ceramide seem to form a similar phase, that is less compatible with POPC. This is remarkable if one thinks about how different these lipids are in their chemical structure, and shows how nature has developed mechanisms to adapt and survive upon defects in the synthesis of particular lipids. This way of interpreting lipid co-regulations, not only in a particular organism or tissue, but also through evolution, explains at least in part the huge lipid diversity present in higher organisms.

Therefore we propose that the common ability of ether PC and ceramide ^44^ to promote lateral segregation, which can be inferred from these experiments with model membranes and the MD simulations, should have consequences for membrane organization, enriching certain regions of the ER membrane with specific lipid species that will help regulate protein function. When either ceramides or EL are depleted from cells, the other lipid amount is adjusted to provide alternative lipids with a similar behavior to carry out important physiological processes, such as membrane traffic in the early secretory pathway. It is thus possible that adaptation of lipid composition is a strategy the cells use to modulate the kinetics of export of proteins from the ER and that EL and SL are at the center of this regulatory hub. In order to examine this, we took advantage of the RUSH system (retention using selective hooks, a technique to synchronize ER export of cargo proteins by addition of biotin ^45^), to investigate the export of GPI-AP under different conditions.

To monitor arrival of secreted proteins to the Golgi we used fluorescent microscopy and quantified the GFP signal from a EGFP-GPI construct in the Golgi, which was segmented from images of cells where the cis-Golgi was stained using an anti-GM130 antibody (Fig 5 A, B, S 3). Due to the heterogeneous behavior within the cell population, we used a high-content automated microscope for image acquisition with which we can measure thousands of cells, which were then filtered to select those in which the fluorescent intensities of the transfected cargo ranged between assigned values, in a consistent way across different conditions (Fig S 3) (see Material and Methods). As a proof of concept, we knocked-down p24 protein (TMED2) and we saw no export of GPI-AP even 10 minutes after the addition of biotin (Fig 5 C-D), showing that TMED2 is required for ER exit of GPI-AP in mammalian cells, which is consistent with the existing literature and with their function as receptors of GPI-AP in yeast for incorporation in COPII carriers ^27, 46, 47^.

**Figure 5.**
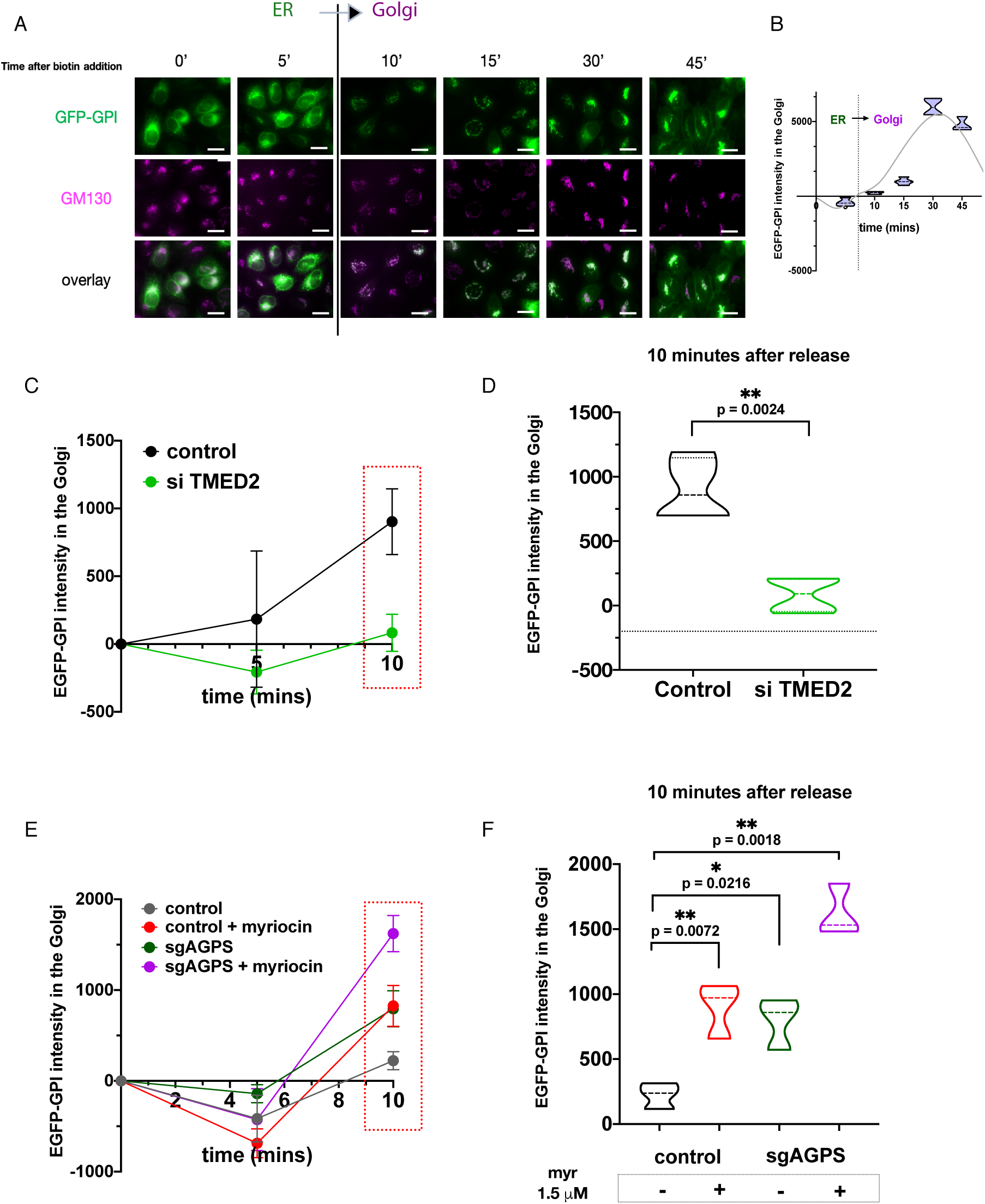
Ether lipid depletion accelerates anterograde transport of GPI-anchored proteins. **a**, Representative images of the RUSH assay in control cells (0, 5, 10, 15, 30 and 45 minutes after release) and p24 knock-down cells (10 minutes after release). The transfected cargo (EGFP-GPI) is released from the ER upon biotin addition and transported to the Golgi apparatus. Scale bars, 20 *µ*m. **b**, Representative quantification of the RUSH assay in control cells 45 minutes after biotin addition. **c**, Kinetics of the export of GPI-anchored proteins upon TMED2 knock-down 10 minutes after biotin addition. **d**, Quantification of the fluorescent intensity of the transfected cargo in the Golgi 10 minutes after biotin addition (control n= 540 cells; TMED2 knock-down n= 1555 cells). **e**, Kinetics of the export of GPI-anchored proteins when cells were treated with myriocin or/and when AGPS was knocked-down. **f**, Quantification of the fluorescent intensity of the transfected cargo in the Golgi 10 minutes after biotin addition (control: n=2198 cells; myriocin: n=1984 cells; sgAGPS: n=2441; sgAGPS + myriocin: n=1398).

When we quantified the arrival of GPI-AP to the Golgi at different time points in SL-depleted and EL-depleted cells we found that 10 minutes after biotin addition, the amount of GFP-GPI localized at the Golgi was higher under those conditions when compared to control cells (Fig 5 E, F), suggesting that these lipid changes caused an acceleration of the transport. Moreover, this effect was cumulative if the cells were subjected to both conditions (SL and EL depletion) at the same time, a situation under which cell viability is compromised (Fig 1 D, G). These experiments measuring ER to Golgi transport of GPI-AP suggest that one of the molecular functions of EL in membranes could be to regulate the selective anterograde transport of GPI-AP (Fig 5 E, F). They also show that regulation of the kinetics of GPI-AP export from the ER is a conserved function of both SL and EL, although the fact that their effects are additive shows that their functions are not completely redundant. In yeast, it has been shown that GPI-AP, which are ceramide based, leave the ER in distinct COPII vesicles with a particular coat protein composition ^48-50^. Similarly, we hypothesize that in mammalian cells the presence of ether lipids as part of the GPI anchor and the selective enrichment of ether PC at these sites, close to p24 proteins could also be a mechanism to facilitate the specific recruitment of GPI-AP and probably separate them from other proteins or lipids that either do not leave the ER, or that do it by carrier vesicles with different protein and lipid composition. As shown above, EL have the ability to segregate from other lipids as has been shown for ceramides.

To examine a putative interaction between EL and the TMD of p24, which is one component of our hypothesis, we performed molecular dynamics (MD) simulations following the protocol previously used by Contreras et al. ^30^. To do so, we tested the interactions of ether PC with p24 TMD in a system containing a mixture of POPC (85%) and ether PC lipids (15%) (Fig 6 A). Interestingly, we observed a selective enrichment of ether PC surrounding the TMD of p24, while the equivalent ester PC (POPC) showed a marked depletion around the protein TM domain (Fig 6 B). The relative abundance of different lipids around the p24 TMD, as measured from their respective radial distribution functions (RDF) showed indeed that ether PC is more often found closer to the TMD of p24 than POPC (Fig 6 C). Of note, when ether PC is present, POPC is depleted from p24 compared to conditions in which POPC is the only lipid in the bilayer (Fig 6 C, black and gray curves). Therefore, another common function of a particular sphingomyelin (C18 n-acyl chain) and ether PC is that both seem to bind to and regulate the function of the p24 complex, which is involved in bidirectional vesicular transport in the early secretory pathway ^30^.

**Figure 6.**
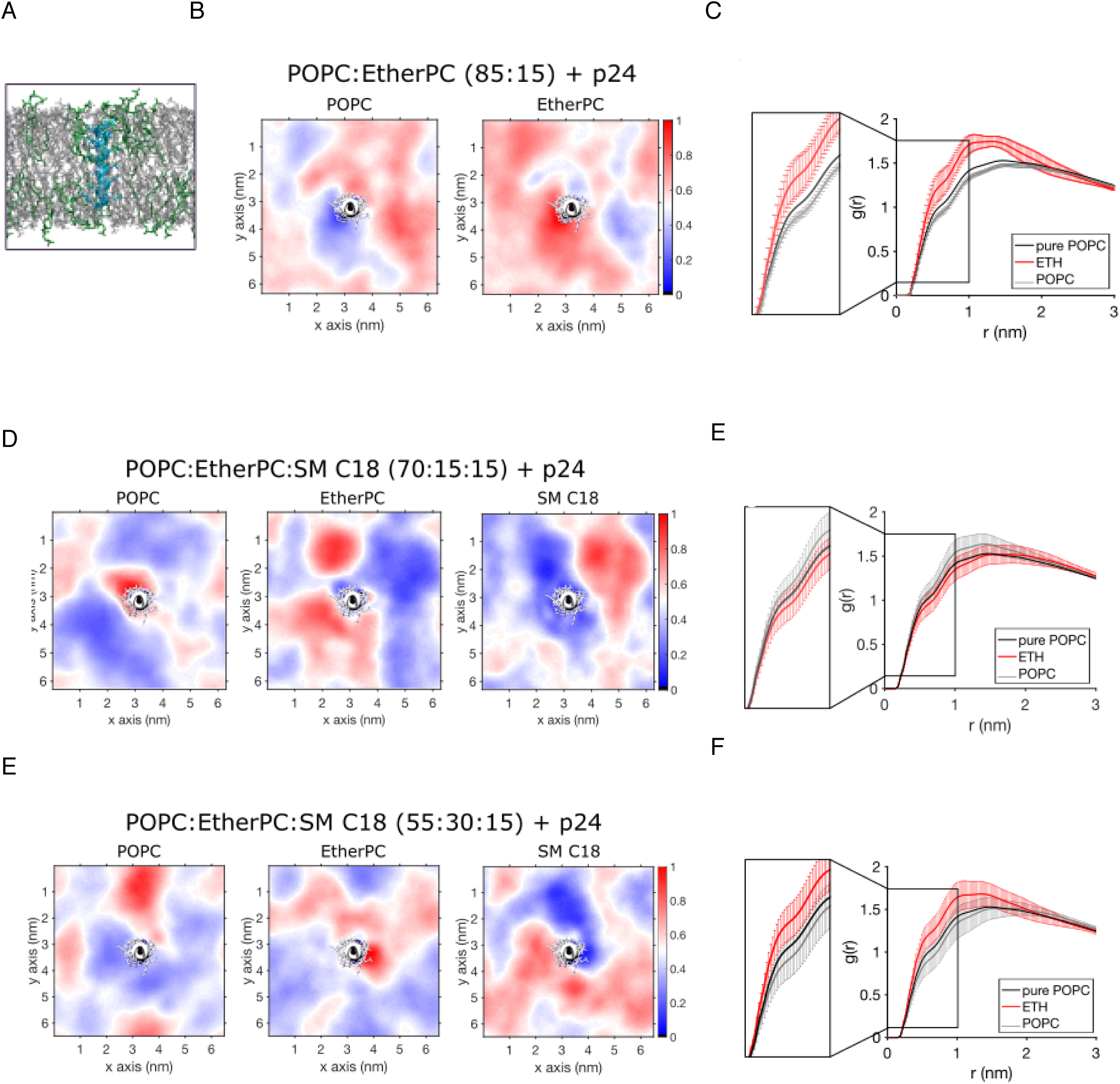
Ether PC can displace POPC from p24 TMD. **a**, Side view of the system POPC (gray):EtherPC (green) 85:15 plus p24 (cyan). **b**, Density maps of POPC and ETH in presence of p24 TMD (white). **c**, Radial distribution function (RDF) of POPC (gray) and ETH (red) with p24 TMD. The curves are compared with the RDF of a system of pure POPC in presence of p24 (black). **d and f**, Density maps of POPC, ETH and SM C18 in presence of p24 (white) at different concentrations. **e and g**, RDF of POPC (gray) and EtherPC (red) with respect to p24 TMD in systems with SM C18 at different molar ratios. The black curve is from a system of pure POPC in presence of p24. In all the images, density maps are obtained as average of the last 500 ns from a representative replica. Red corresponds to enrichment, while blue corresponds to depletion of the lipids. For better clarity, only RDF of Ether PC and POPC are displayed. Please see Figure S4 B and C for the curves of SM C18.

In order to characterize the possible competition between SM C18 and ether PC for the binding of the p24 TMD, we investigated the behavior of a system consisting of POPC, SM C18 and p24 TMD. There, we could successfully reproduce the previously described binding of SM C18 to the p24 TMD (Fig S4 A, E). Next, we analyzed bilayers composed of a ternary mixture of POPC, ether PC and SM C18 at two different lipid ratios: one in which ether PC and SM were present in the same amount and another one in which ether PC was twice the amount of SM. Our results indicate that when ether PC and SM C18 are present at equal concentrations, the two lipids compete for the space close to p24, resulting in a displacement of the ether PC from the surroundings of p24 (Fig 6 D, E, S4 B). In contrast, when ether PC is present at a higher concentration than SM C18, a condition that is more likely to exist under physiological conditions due to the relative amounts of these two lipids in cells, ether PC is able to surround the p24 TMD, pushing away POPC and SM C18 (Fig 6 F, G, S4 C). Taken together, our simulations suggest that ether PC and SM C18 interact with p24 using different mechanisms; while ether PC molecules are more stably associated with the p24 TMD, only few SM molecules interact with p24 at once and they display a very dynamic behavior. As a consequence, SM C18 can be found at either very high or very low concentrations close to p24, resulting in large error bars in radial distribution functions obtained from different replicas (Fig S4 B and C). Our results, while consistent with the mechanism for SM binding to p24 previously shown, where only one molecule of SM C18 interacts with the SL binding domain of p24 at a given time, strongly suggest a different role for ether PC when compared to SM18. SM18 binds to selected residues of the p24 TMD while ether PC provides a more structural and scaffolding role around the protein. Different binding mechanisms are compatible with the proposed model and the dual function of p24 proteins in the secretory pathway, as EL would help to have selective anterograde transport of GPI-AP, while SL would be required for their COPI-dependent retrograde transport back to the ER. Altogether, the maintenance of the proper levels of these two lipid classes would help maintain the homeostasis of the p24 protein cycle.

In summary, we have uncovered a genetic interaction and a metabolic co-regulation of EL and SL that suggest common functions for these two lipid classes. Further validation using *in vitro* experiments and MD simulations shows how ceramide and ether PC, although having very different structures, display similar effects on membrane organization and share the ability to segregate in membranes. In addition, we find multiple lines of evidence pointing to a specific role of both lipid classes in controlling p24-mediated transport in the early secretory pathway. First, knockdown of p24 proteins (TMED2, TMED10) renders cells hypersensitive to SL depletion. Second, lipidomic analysis of EL-deficient cells shows a concurrent decrease in SM C18 levels, which is known to specifically regulate p24 ER-to-Golgi cycling. Third, high-content live-cell microscopy shows that depletion of both SL and EL impact transport of GPI-AP between ER and Golgi. Finally, MD simulations validate a molecular cross-talk between both lipid classes and p24 TMD. Together, our data highlights a new role for lipids as direct regulators of intracellular transport, and suggest that the lipid diversity found in eukaryotic cells exists in part to ensure the homeostasis of the secretory pathway.

### Evolutionary implications

The identification of new molecular functions of EL has important implications for the field of membrane evolution. Our results confirm that even though major aspects of the secretory pathway are conserved between yeast and human, some regulatory components like lipids, and as a consequence, lipid-protein interactions, have been modified and have undergone convergent evolution to meet the needs of each organism. EL are very abundant lipids in mammalian cells although they seem to be absent from fungi and plants. In yeast, Cer is used for the remodeling of the GPI anchor while EL have been selected to perform this function in mammalian cells. We propose that the basis for this selection is related to the shared physico-chemical aspects of these lipids and the effects they exert on membranes. Our results show how the discovery of co-regulated lipids species across the evolutionary tree can improve our understanding of their molecular functions.

## MATERIALS AND METHODS

### Cell lines, culture conditions, reagents and antibodies

K562 cells were purchased from ATCC (ATCC^®^ HTB-56™) and cultured as shown in the section regarding the CRISPRi screen. HeLa (Kyoto) cells were a gift from Anthony Hyman group (MPI-CBG) and were cultured at 37°C and 5 % CO_2_ in Dulbecco’s Modified Eagle Medium (Cat# 31966047, DMEM, GIBCO™) with 4.5 g/L glucose, supplemented with 10 % delipidated fetal calf serum (FCS, Hyclone) and 1 % Pen/Strep (Cat# 10378016, GIBCO™). BV2 cells were provided by Prof. Thierry Soldati (University of Geneva) and Calu-6 cells were purchased from ATCC (ATCC^®^ HTB-56™). Both cell lines were cultured in the same conditions as Hela cells. The number of cells was determined using a Countess II FL Automated Cell Counter and Countess™ Cell Counting Chamber Slides which include the trypan blue solution (Thermo Fisher Scientific). Transfections with siRNAs were performed using Lipofectamine RNAiMax (Cat# 13778075, Thermo Fisher Scientific) using the supplier’s instructions, for 3 days, if other conditions are not specified. The siRNA solutions were diluted using Opti-MEM® (Cat# 31985047). The siRNAs used in this work were: siPOOL-AGPS (Gene ID: 8540), siPOOL-TMED2 (Gene ID: 10959), siPOOL-SMPD1 (Gene ID: 6609), siPOOL-SMPD3 (Gene ID: 55512), and siPOOL non-coding (nc) siRNA (siTOOLs BIOTECH, Planegg, Germany). *Trans*IT-X2® Dynamic Delivery System (Mirus Bio) was used to transfect plasmid DNA. Str-KDEL_SBP-EGFP-GPI construct was a gift from Franck Perez (Addgene plasmid #65294).

#### Serum de-lipidation

Serum de-lipidation was achieved using lipid removal adsorbent (LRA, MilliporeSigma #13358-U). 500mL of fetal bovine serum (FBS, HyClone GE Life Sciences) was incubated overnight with 25g LRA resin under constant rotation at 4C. Resin particles were then decanted by centrifugation (first spin for 5min at 2,000xg for bulk removal, second spin at 27,000xg for fine particle removal). FBS pH was adjusted to 7.4 with NaOH and filter-sterilized using a 0.2*µ*m membrane.

### Antibodies

Purified mouse anti GM130 (Cat# 610822, BD Biosciences). Alexa Fluor® 647-AffiniPure Donkey Anti-Mouse IgG (H+L) (Cat# 715-605-150, Jackson ImmunoResearch).

### MTT assay

The MTT(3-[4,5-dimethylthiazol-2-yl]-2,5-diphenyltetrazolium bromide; thiazolyl blue) (Cat# M6494, Thermo Fisher Scientific) assays were performed in 96-well black/clear bottom plates (FALCON®, Corning). MTT was dissolved in Dulbecco’s Modified Eagle Medium without phenol red to a 5 mg/ml stock solution, added to each well to be one-tenth the original culture volume and incubated for 3 to 4 hr. At the end of the incubation period the medium was removed and the dye was solubilized with DMSO : isopropanol (1:1). Absorbance of converted dye is measured at a wavelength of 570 nm with background subtraction at 630–690 nm.

#### CRISPRi screen

Screens were performed in K562 cells stably expressing a constitutive SFFV-dCas9-BFP-KRAB CRISPRi transcriptional repression construct using lentiviral transduction ^51^, Addgene plasmid # 46911). K562 CRISPRi cells were grown in RPMI 1640 Hepes medium supplemented with 10% lipid-depleted FBS and transduced with a genome-wide sgRNA library targeting the transcriptional start site of ∼16,000 protein-coding genes as well as 1,000 negative control scrambled sgRNA sequences as in ^32^(Addgene pooled library # 62217). Each protein-coding gene was targeted by a minimum of 10 unique sgRNA sequences to insure statistical power, and cell culture was scaled to insure a minimum representation of 1,000 cells per unique sgRNA at all times. sgRNA-expressing cells were puromycin-selected as described ^32^ and a “time zero” sample was isolated and the cell pool was subsequently split into two parallel 1-liter cultures in spinner flasks under constant agitation. One culture was treated with 1*µ*M myriocin (MilliporeSigma #M1177, resuspended a 1mM in DMSO) while a control culture was treated with DMSO. Sphingolipid depletion was insured by myriocin treatment for 6 days, followed by 6 days of recovery in drug-free media. Cells from both cultures were then harvested for bulk genomic DNA extraction, and the sgRNA-encoding regions were then amplified by PCR and sequenced on an Illumina HiSeq-2500 as described ^32^. Myriocin-specific phenotypes (ρ) were quantified by comparing the frequency of cells harboring a given sgRNA sequence between the treated and un-treated cultures, while growth phenotypes (γ) were quantified by comparing, within each culture, sgRNA frequencies between the final cell pools and the “time zero” sample ^32, 33^. Bioinformatic analysis was performed as described ^32^. Briefly, hit genes were scored based on the average phenotype of the 3 strongest sgRNAs (by absolute value) targeting their TSS using a Mann-Whitney test against the non-targeting control set. For each gene, the myriocin-specific gene phenotype representing resistance to treatment (ρ) was calculated as described ^33^ and forms the basis of our analysis. Both growth (γ) and myriocin-specific (ρ) phenotypes for each genes are included in the Supplementary file Gene_phenotype_table.xslx.

### Lipids

All the lipids purchased for this study were from Avanti Polar Lipids, Inc. (Alabaster, AL): Cer 24:1 (Cat# 860525), POPC (Cat# 850457). Myriocin from *Mircelia Sterilia* was purchased from Sigma-Aldrich-Fluka (Cat# M1177).

#### Synthesis of ether PC

To a solution of Lyso-PAF C16 (10.9 mg, 0.023 mmol) in chloroform (1.0 mL), oleoyl chloride (30 μL), DMAP (cat. amount), and DIPEA (40 μL) was added. The reaction mixture was stirred at 55 °C for 2 hours. The crude product was directly purified by flash chromatograph (SiO_2_, first MeOH/DCM (1/10), then MeOH/DCM/H_2_O (2/1/0.1, v/v)) to yield the product as a solid (13.5 mg, yield: 77 %). The reaction and purification were monitored by thin layer chromatograph (TLC) using phosphomolybdic acid (PMA) staining.

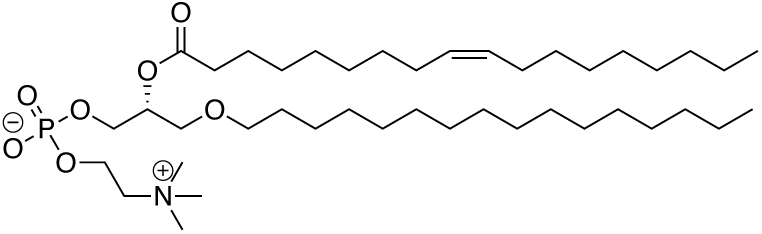

### Lipid extraction

For lipidomics experiments 3×10^5^ cells were seeded in 6 cm dishes and harvested after 3 days of culture in delipidated medium. Lipid extraction was performed using a modified MTBE protocol ^52^. Cells were first washed with cold PBS and scraped off in 500 µl cold PBS on ice. The suspension was transferred to a 2 ml Eppendorf tube and spin down at 3200 rpm for 5 minutes at 4°C. After removing the PBS, samples were stored at -20°C or directly used for further extraction. Then, 360 μl methanol was added and vortexed. A mixture of lipid standards (see Key Resources Table) was added and the cells were vortexed for 10 minutes at 4°C using a Cell Disruptor Genie (Scientific Industries, Inc). MTBE (1.2 mL) was then added and the samples were incubated for one hour at room temperature with shaking (750 rpm). Phase separation was induced by adding 200 μl H_2_O. After 10 min of incubation at RT, the sample was centrifuged at 1000 x g for 10 min (RT). The upper (organic) phase was transferred in a 13 mm screw cap glass tube and the lower phase was extracted with 400 μl artificial upper phase (MTBE/methanol/water (10:3:1.5, v/v)). The two upper phases were combined and the total lipid extract was divided in 3 equal aliquots (one for phospholipids (TL), one for sterols (S) in 2 mL amber vials and one for sphingolipids ^53^ in a 13 mm glass tube) and dried in a Centrivap at 50°C or under a nitrogen flow. The SL aliquot was deacylated to eliminate phospholipids by methylamine treatment (Clarke method). 0.5 mL monomethylamine reagent (MeOH/H2O/n-butanol/Methylamine solution (4:3:1:5 v/v) was added to the dried lipid, followed by sonication (5 min). Samples were then mixed and incubated for one hour at 53°C and dried (as above). The monomethylamine treated lipids were desalted by n-butanol extraction. 300 μl H_2_O saturated n-butanol was added to the dried lipids. The sample was vortexed, sonicated for 5 min and 150 μl MS grade water was added. The mixture was vortexed thoroughly and centrifuged at 3200 x g for 10 min. The upper phase was transferred in a 2 mL amber vial. The lower phase was extracted twice more with 300 μl H_2_O saturated n-butanol and the upper phases were combined and dried (as above).

### Glycerophospholipid and Sphingolipid Detection on a Triple Quadrupole TSQ Vantage (ThermoFischer Scientific)

TL and SL aliquots were resuspended in 250 μl Chloroform/methanol (1:1 v/v) (LC-MS/HPLC GRADE) and sonicated for 5 minutes. The samples were pipetted in a 96 well plate (final volume = 100 μl). The TL were diluted 1:4 in negative mode solvent (Chloroform/Methanol (1:2) + 5mM Ammonium acetate) and 1:10 in positive mode solvent (Chloroform/Methanol/Water (2:7:1 v/v) + 5mM Ammonium Acetate). The SL were diluted 1:10 in positive mode solvent and infused onto the mass spectrometer. Tandem mass spectrometry for the identification and quantification of lipid molecular species was performed using Multiple Reaction Monitoring (MRM) with a TSQ Vantage Triple Stage Quadrupole Mass Spectrometer (Thermo Fisher Scientific) equipped with a robotic nanoflow ion source, Nanomate HD (Advion Biosciences, Ithaca, NY). The collision energy was optimized for each lipid class. The detection conditions for each lipid class are listed below. Ceramide species were also quantified with a loss of water in the first quadrupole. Each biological replicate was read in 2 technical replicates (TR). Each TR comprised 3 measurements for each transition. Lipid concentrations were calculated relative to the relevant internal standards and then normalized to the total lipid content of each lipid extract (mol%).

### Automated fluorescence light microscopy

For the high content microscopy experiments we used black 96-well imaging plates (cat# 89626, Ibidi). 3000 cells were seeded per well in media containing either the vehicle (MetOH) or 1.5 *µ*M myriocin from a 5 mM stock solution. A reverse transfection has been performed for each siRNA experiment, using lipofectamine RNAimax following manufacturer instructions. After 48 hours of incubation, DNA transfection was performed for another 24 hours. Briefly, the medium from each well was replaced with 120 µL of fresh media added to a 30 µL DNA solution previously diluted in Opti-MEM and mixed with *Trans*IT-X2® Dynamic Delivery System reagent in a 3:1 *Trans*IT-X2 (µL): DNA (µg) ratio.

#### Retention using selective hooks (RUSH)

Before the experiment was performed, a solution of biotin was prepared in media at a concentration of 120 *µ*M (40 *µ*M final concentration in each well). Media was replaced in each well by 100 µL of fresh media and the cells were kept in the incubator for at least 30 minutes before the experiment. To start the release of the transfected fluorescent cargo, 50 µL of the biotin solution were added at different time points (45, 30, 15, 10, and 5 minutes). After 45 minutes the plate was fixed using 4% PFA solution and washed using an automated plate washer (BioTek EL406). Cells were then stained as follows: step 1: monoclonal antibody against GM130, 1/500, saponin 0.05%, BSA 1%, in PBS, incubation for 1 h and, step 2: Hoechst 33342 Solution (20 mM) (1/5000), Cy5-labelled secondary antibody against mouse IgG, 1/500, incubation for 30 min. Image acquisition was performed immediately after staining using a ImageXpress® Micro Confocal High-Content Imaging System (Molecular devices) with the 40× objective. 36 images were captured per well. For image analysis, we used the MetaXpress Custom Module editor software to first segment the image and generate relevant masks (Fig S3). In the first step, individual cells were identified using the staining of the nuclei (Hoechst channel). Next, the Golgi was segmented from the images using the signal coming from the anti-GM130 antibody (Cy5 channel, Fig S3). Properly transfected cells were then selected using those ones with a fluorescence intensity of the cargo (EGFP-GPI) ranging between two specific values, identical through all conditions. Finally, the masks were applied to the original fluorescent images, and different measurements were obtained per cell (e.g. integrated intensity, average intensity and object count). The average intensity value of the fluorescent cargo in the Golgi is used to represent the data. The same imaging and analysis pipelines were applied to all images. Data analysis was with Prism Graph Pad 8.0.

### Giant unilamellar vesicle (GUV) formation and imaging

Giant unilamellar vesicles (GUVs) were prepared by the agarose method ^54^. Stock lipid solutions (2 mg/ml total lipid) were prepared in chloroform: methanol (2:1, v/v) and spread on ultra low gelling agarose (Sigma-Aldrich # A2576) (1%)-coated glass slides. First, the glass slides were dipped in the agarose solution and dried by keeping the slides for 2 h at 30°C or overnight at room temperature. Then the lipid mixtures were spread onto the agarose-coated slides. For GUV formation, a solution of 280mM sucrose was added to each slide and the chambers were placed at the desired temperature, always above the transition temperature of the lipid mixtures. For the confocal microscopy: a home-made chamber was used that allows direct GUV visualization under the microscope ^55, 56^.

### Fluorescence lifetime measurements

FLIM imaging was performed using a Nikon Eclipse Ti A1R microscope equipped with a Time Correlated Single-Photon Counting module from PicoQuant as previously published ^58, 59^. Excitation was performed using a pulsed 485nm laser (PicoQuant, LDH-D-C-485) operating at 20 MHz, and emission signal was collected through a bandpass 600/50nm filter using a gated PMA hybrid 40 detector and a TimeHarp 260 PICO board (PicoQuant). SymPhoTime 64 software (PicoQuant) was then used to fit fluorescence decay data (from full images or regions of interest) to a dual exponential model after deconvolution for the instrument response function (measured using the backscattered emission light of a 1μM fluorescein solution with 4M KI). Data was expressed as means ± standard deviation of the mean. The full width at half-maximum (FWHM) response of the instrument was measured at 176 ps.

### Molecular dynamic simulations

Molecular dynamics simulations were run with the GROMACS ^60^ software using the CHARMM36 force field ^61, 62^. The coordinates of p24 TMD were used as described ^30^. The CHARMM GUI *Membrane Builder* ^*63-65*^ was used to build systems containing p24 already inserted in the bilayer, 102 lipids, 50 water/molecules per lipid and an ion concentration of 150 mmol/L. The same approach was used to build pure lipid bilayers, with 100 lipids per leaflet and 70 water molecules per lipid. For the systems containing ether PC, as this lipid is not natively present in CHARMM-GUI, lipid bilayers containing SOPC were built and then atom types were modified with an in-house script for conversion to ether PC.

The simulations containing p24 were equilibrated using the CHARMM GUI procedure that involves minimization followed by steps in which positional restraints for lipid head groups and protein backbone are gradually lowered and removed. The systems were minimized using a steepest descent algorithm until reaching a force smaller than 1000 kJ mol^-1^ nm^-1^. A 175000 step equilibration with a time step increasing from 0.001 to 0.002 ps was run, followed by a 1000 ns production, with a time step of 0.002 ps. In both equilibration and production, pressure was controlled by a Parrinello-Rahman barostat ^66^ and average pressure was 1 atm, coupled every 5 ps. During equilibration, a Berendsen thermostat ^67^ was used, while during production temperature was controlled via a Nosé-Hoover thermostat ^68^, coupled every 1 ps and the target temperature was 310K during the entire run. The lateral *xy* dimensions were coupled, while the *z* dimension was allowed to fluctuate independently. Van der Waals and electrostatic interactions were truncated at 1.2 nm. Fast smooth Particle-Mesh Ewald (SPME) ^69^ was used for calculations of electrostatics. For Van der Waals calculations a twin range cut-off with neighbor list cut-off coupled with a function to smoothly switch the potential to 0 from 1 nm was used.

For systems containing only lipid bilayers the same settings were used, but with a different equilibration protocol: a minimization using a steepest descent algorithm until reaching a force smaller than 700 kJ mol^-1^ nm^-1^ was performed, followed by a 20 ps equilibration with a time step of 0.001 and using a Nosé-Hoover thermostat for temperature control. Finally, productions were run for 200 ns, with a time step of 0.002 ps. Two independent replicas for each lipid compositions (see table 2) were run.

**Table 1:**
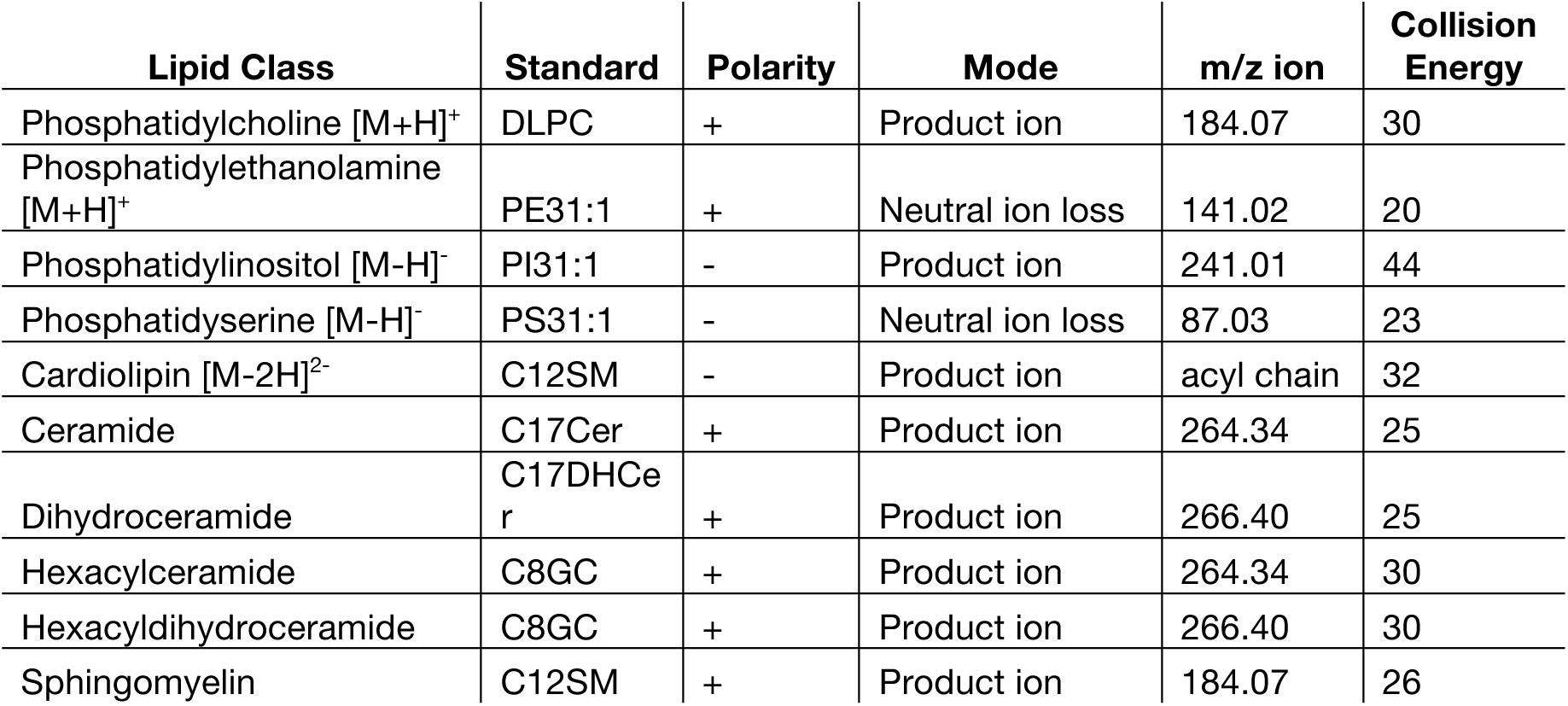
Detection of Lipids by MS/MS (Related to Lipidome Analyses)

| Lipid Class | Standard | Polarity | Mode | m/z ion | Collision Energy |
| --- | --- | --- | --- | --- | --- |
| Phosphatidylcholine [M+H]<sup>+</sup> | DLPC | + | Product ion | 184.07 | 30 |
| Phosphatidylethanolamine [M+H]<sup>+</sup> | PE31:1 | + | Neutral ion loss | 141.02 | 20 |
| Phosphatidylinositol [M-H]<sup>-</sup> | PI31:1 | - | Product ion | 241.01 | 44 |
| Phosphatidylserine [M-H]<sup>-</sup> | PS31:1 | - | Neutral ion loss | 87.03 | 23 |
| Cardiolipin [M-2H]<sup>2-</sup> | C12SM | - | Product ion | acyl chain | 32 |
| Ceramide | C17Cer | + | Product ion | 264.34 | 25 |
| Dihydroceramide | C17DHCer | + | Product ion | 266.40 | 25 |
| Hexacylceramide | C8GC | + | Product ion | 264.34 | 30 |
| Hexacyldihydroceramide | C8GC | + | Product ion | 266.40 | 30 |
| Sphingomyelin | C12SM | + | Product ion | 184.07 | 26 |

**Table 2:** List of all the MD setups, with bilayer composition, number of replicas and length of simulations.

| Composition | Ratio | Number of replicas | Length (ns) |
| --- | --- | --- | --- |
| POPC: Cer | 80:20 | 2 | 200 |
| ether PC: Cer | 80:20 | 2 | 200 |
| POPC: ether PC: Cer | 40:40:20 | 2 | 200 |
| POPC: ether PC | 85:15 | 2 | 200 |
| POPC: ether PC | 70:30 | 2 | 200 |
| POPC + p24 | 100 | 2 | 1000 |
| POPC: SM C18 + p24 | 85:15 | 2 | 1000 |
| POPC: ether PC + p24 | 85:15 | 8 | 1000 |
| POPC: ether PC: SM C18+ p24 | 70:15:15 | 8 | 1000 |
| POPC: ether PC: SM C18+ p24 | 55:30:15 | 8 | 1000 |

For the systems containing both ether PC and p24, for each different lipid composition, 8 independent 1 μs-long replicas were run. For the systems “POPC + p24” and “POPC:SM C18 + p24”, 2 independent 1 μs-long replicas were run. The last 500 ns were used for analysis. From radial distribution function (RDF) calculations, obtained using a VMD ^70^ plugin, it appeared that some systems were not converged. In order to select only the converged simulations for the final analysis, the RDF curves of ether PC from all the 8 replicas were averaged and similarity between the RDFs of each single replica and the average curve was calculated through the “Frechet Distance Calculator” script for MATLAB (Figure S5). Only curves with a similarity below a cutoff of 0.2 were selected and used to calculate the average RDFs and relative standard deviation shown in Fig 3 A, C and E.

For the systems with pure lipid bilayers, 2 independent 200 ns-long replicas were run and the last 100 ns were used for analysis.

Density maps for the systems containing p24 were calculated using GROMACS tools, as average of the last 500 ns of a representative replica per system. To obtain an enrichment/depletion measurement, the density maps of the single lipids were normalized at the same relative concentration. Then, the normalized density maps were summed and the density maps of the protein was subtracted (sum-densmap). Finally, the normalized density map of each lipid was divided by the sum-densmap. For the systems with only lipids, density maps were obtained from the last 100 ns of simulations.

Hydrogen bonds calculation was performed using GROMACS tools.

### RNA extraction and qPCR

Cells were grown in 6 cm dishes. Total RNA extraction was performed using the RNeasy Mini Kit (Qiagen No: 74104). RNA quantity and purity were controlled using a spectrophotometer: ratios 260/280 (protein contamination) and 260/230 (guanidinium thiocyanate contamination) were greater than or equal to 2.0.

One microgram of RNA was used for cDNA synthesis using the Superscript II (Invitrogen) reverse transcription reaction with random hexamers. qPCR analysis was performed on a CFX Connect Real-Time PCR system using SsoAdvanced Universal SYBR Green supermix (Bio-Rad, Switzerland).

Primers were designed using Primer-BLAST, and sequences were the following (5’->3’):

AGPS-forward: GAGTGCAAAGCGCGGAGA.

AGPS reverse: TTCTTGCCGCTTCTTTGGGA.

TATA-binding protein (TBP)-forward: CCGCCGGCTGTTTAACTTC.

TATA-binding protein-reverse: AGAAACAGTGATGCTGGGTCA.

The amplification profile for qPCR was 95°C, 1 min, followed by 40 amplification cycles at 95°C, 10 s, and 60°C, 30 s. The run was completed with the dissociation curve from the start at 65°C to 95°C with 0.5 increments. Data analysis was done with the Bio-Rad CFX manager software version 3.1. Expression of the target gene was calculated relative to the wild-type condition after normalization to the reference gene TBP. Triple biological replicates and triple technical replicates were analyzed.

## QUANTIFICATION AND STATISTICAL ANALYSIS

Statistical analyses and data plotting were performed using Prism Graph Pad 8.0. An unpaired, two tailed, unequal variance t-test assuming Gaussian distribution was used to compare between two groups. Data represent mean ± SEM and p-values are shown in the figures.

## ACKNOWLEDGEMENTS

NJR was supported by a Postdoctoral fellowship from the Basque Government. Manuel D. Leonetti was supported by a HHMI fellowship from the Jane Coffin Childs Memorial Fund for Medical Research. VZ and SV acknowledge support by the Swiss National Science Foundation (grant #163966). This work was supported by grants from the Swiss National Supercomputing Centre (CSCS) under project ID s726 and s842. HR is supported by the Swiss National Science Foundation and the NCCR Chemical Biology (grants 183561, 184949, 185898). We thank Isabelle Riezman for technical support in mass spectrometry and Dimitri Moreau and Stefania Vossio from the ACCESS Geneva High-content microscopy facility for support in microscopy and data analysis.

## AUTHOR INFORMATION

These authors contributed equally: Noemi Jiménez-Rojo and Manuel D. Leonetti.

### Author contributions

NJR, MDL, HR and JSW designed the work. NJR performed most of the experiments with input from HR, SV, MDL, AC and AR. MDL performed the CRISPRi screen. VZ and SV performed and analyzed the MD simulations. AC and AR performed and/or assisted with the biophysical experiments. NRI assisted with the first RUSH measurements. SM provided the FLIPPER-TR®. SF synthesized the ether PC. NJR and HR wrote the manuscript with input from co-authors.

## ETHICS DECLARATIONS

### Competing interests

The authors declare no competing interests.

