## Supplemental figures for "Conserved function of ether lipids and sphingolipids in the early secretory pathway"

Figure S1.

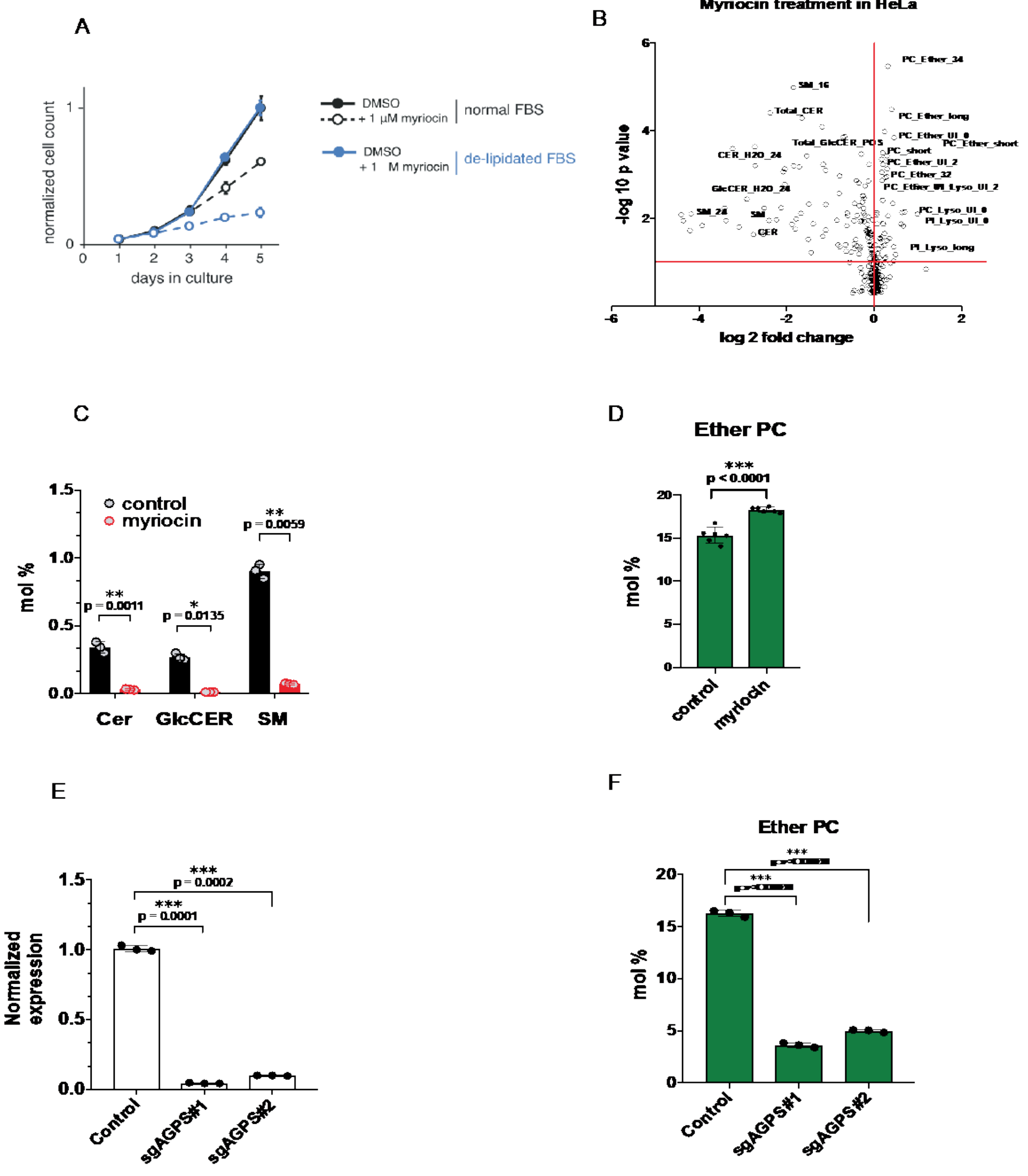


**Supplementary Figure 1.**

**a,** Growth curves are shown for K562 cells grown in media containing normal or de-lipidated fetal bovine serum (FBS), and treated with 1 µM myriocin for the indicated time period. In both normal and delipidated conditions, cell counts are normalized to day 5 counts for respective untreated controls. Each point represents average ± standard deviation from 3 independent measurements. **b,** Lipid changes in Hela cells after 3-day inhibition of sphingolipid synthesis with 1.5 µM myriocin. Log 2 of the fold change to control cells in the X-axis and the -log P value in the Y-axis (n=3). **c,** Relative sphingolipid levels over the total of lipids detected and after 1.5 µM myriocin treatment in Hela cells (n=3). **d,** Relative ether phosphatidylcholine (PC) levels upon myriocin treatment (n=6). **e,** Knockdown efficiency of two different sgRNA targeting AGPS and measured by quantitative real-time PCR (qRT-PCR) (n=3). **f,** Relative ether PC levels over the total of lipids detected in the two different AGPS CRISPRi cell lines (n=3).

Figure S2.

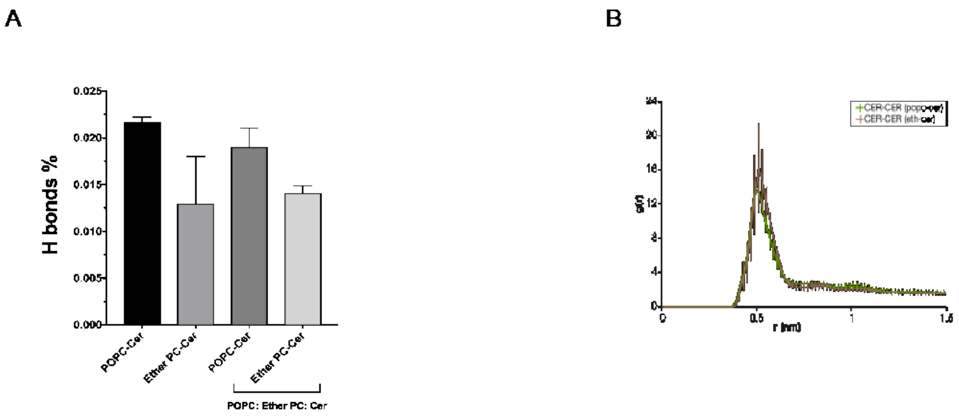


**Supplementary Figure 2.**

**a,** Percentage of hydrogen bonds between POPC and Cer or Ether PC and Cer in different systems calculated from MD simulations. **b,** Radial distribution function of Cer-Cer in binary mixtures with POPC or Ether PC from MD simulations

Figure S3

### Retention using selective hooks (RUSH): image analysis pipeline

**1. Cell body determination:** the Hoechst 33342 channel is used to identify individual cells

**2. Golgi segmentation:** The Golgi is segmented after applying a top hat image modification (GM130, Cy5 channel) to lower the background signal and facilitate the segmentation.

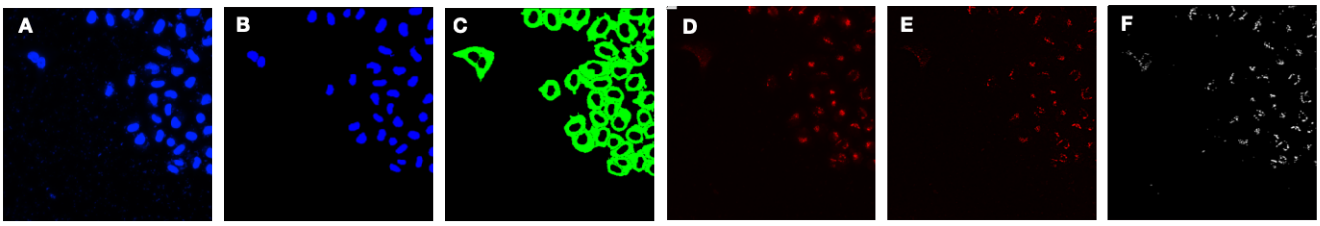


**3. Selection of properly transfected cells:** a filtering, based on a range of intensities in the GFP channel, is used to positively selected the transfected cells. Those are the only cells kept for further quantification.

**4. Quantification of fluorescent cargo intensity in the Golgi per cell:** the GFP intensity (average and integrated) in the Golgi mask is extracted for each positive cell. An average over all images in a well is used for the plotting.

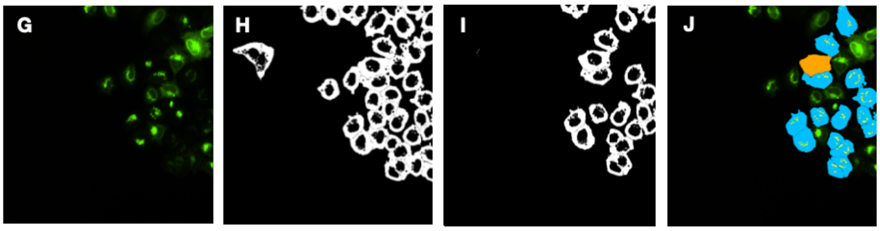


| ObjectID | Integrated Intesity | Average Intensity |
|---|---|---|
| 1 | 20703076 | 16419.54 |
| 2 | 2571141 | 2394.32 |
| 3 | 24884288 | 14243.64 |
| 4 | 12790525 | 6848.19 |

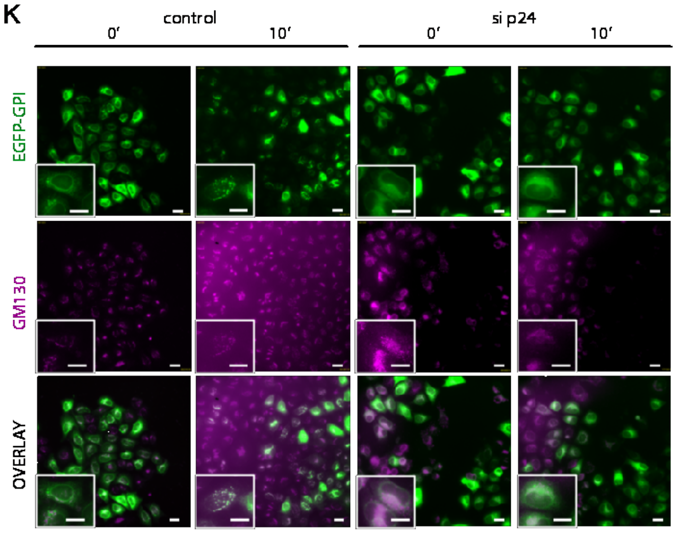


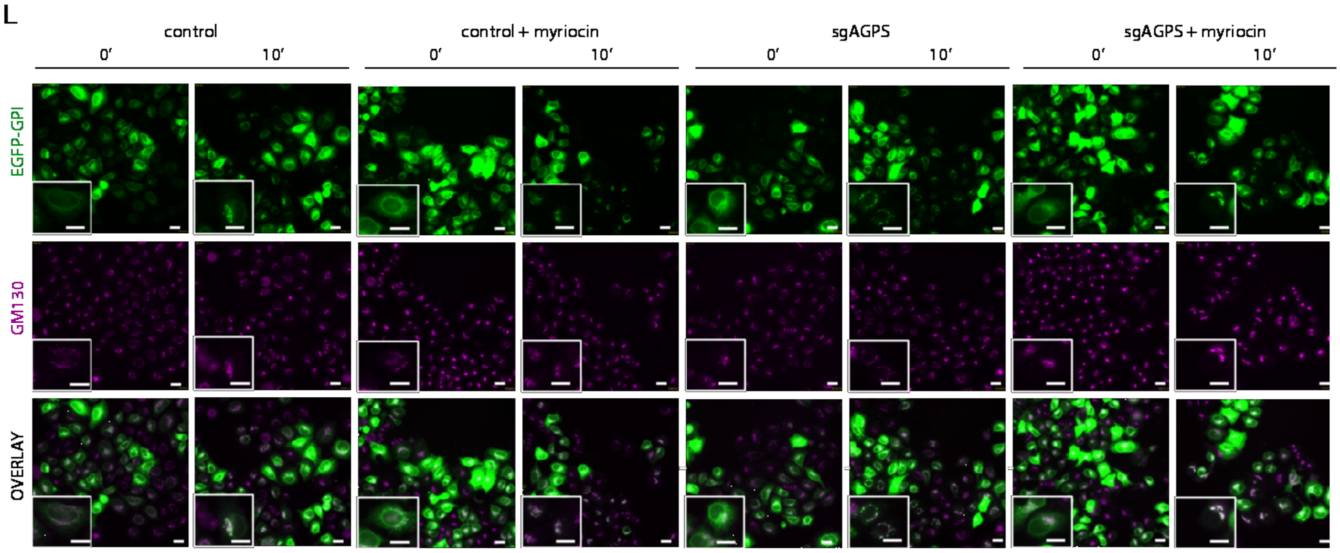

**Supplementary Figure 3. Image analysis pipeline for the RUSH data (retention using selective hooks).**

**a,** Original image, Hoechst channel. **b,** Nuclei mask. **c,** Cell mask. **d,** Original image, Cy5 channel (Golgi). **e,** Image after applying a top hat modification to remove the background, Cy5 channel. **f,** Golgi mask, Cy5 channel. **g,** Original image, GFP channel (EGFP-GPI). **h,** Cytoplasm mask. **i,** Cytoplasm mask of selected cells for further analysis**. j,** Original image, GFP channel, with the representative mask and a table with intensity values per cell in the image. **k,** Representative microscopy images of the RUSH experiment in control Hela cells and cells where TMED2 was knocked-down. 0'=before biotin addition; 10'= 10 minutes after biotin addition). **l,** Representative microscopy images of the RUSH experiment in control Hela cells, cells where AGPS was knocked-down, cells treated with myriocin, and cells where both treatments are combined. 0'=before biotin addition; 10'= 10 minutes after biotin addition).

Figure S4.

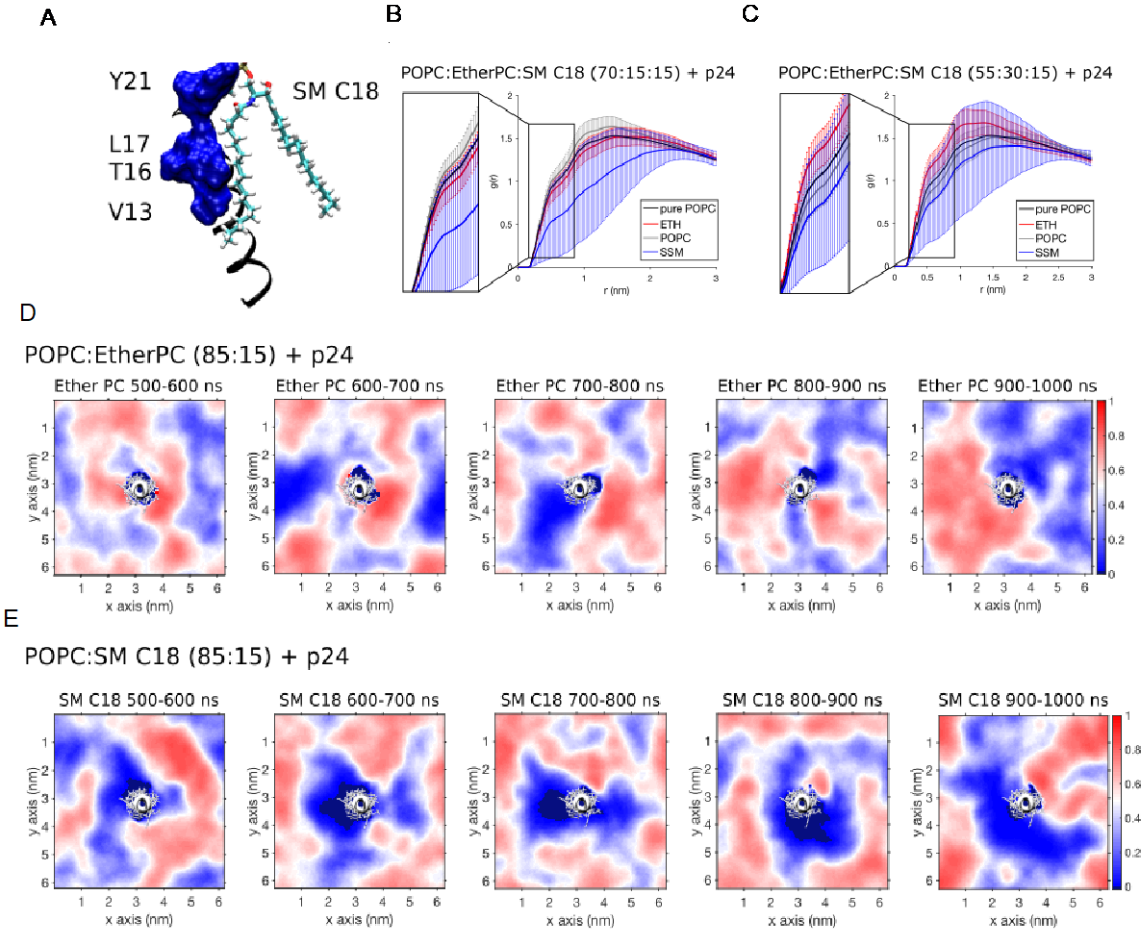


**Supplementary Figure 4. Ether PC and SM C18 display different binding mechanisms to p24 TMD.**

**a,** Binding of SM C18 to the sphingolipid binding domain (V13, T16, L17, Y21) of p24 TMD, as in Contreras et al. **b and c**, RDF of POPC (gray), Ether PC (red) and SM C18 (blue) with respect to p24 TMD in systems with SM C18 at different molar ratios. The black curve is from a system of pure POPC in presence of p24. **d,** Density maps of the distribution of ether PC in the last 500 ns of simulations, averaging every 100 ns, from a representative replica. **e,** Density maps of the distribution of SM C18 in the last 500 ns of simulations, averaging every 100 ns, from a representative replica. While ether PC is stably close to p24 TMD, SM C18 interacts with a small portion of the protein with an on/off mechanism.

.

Figure S5.

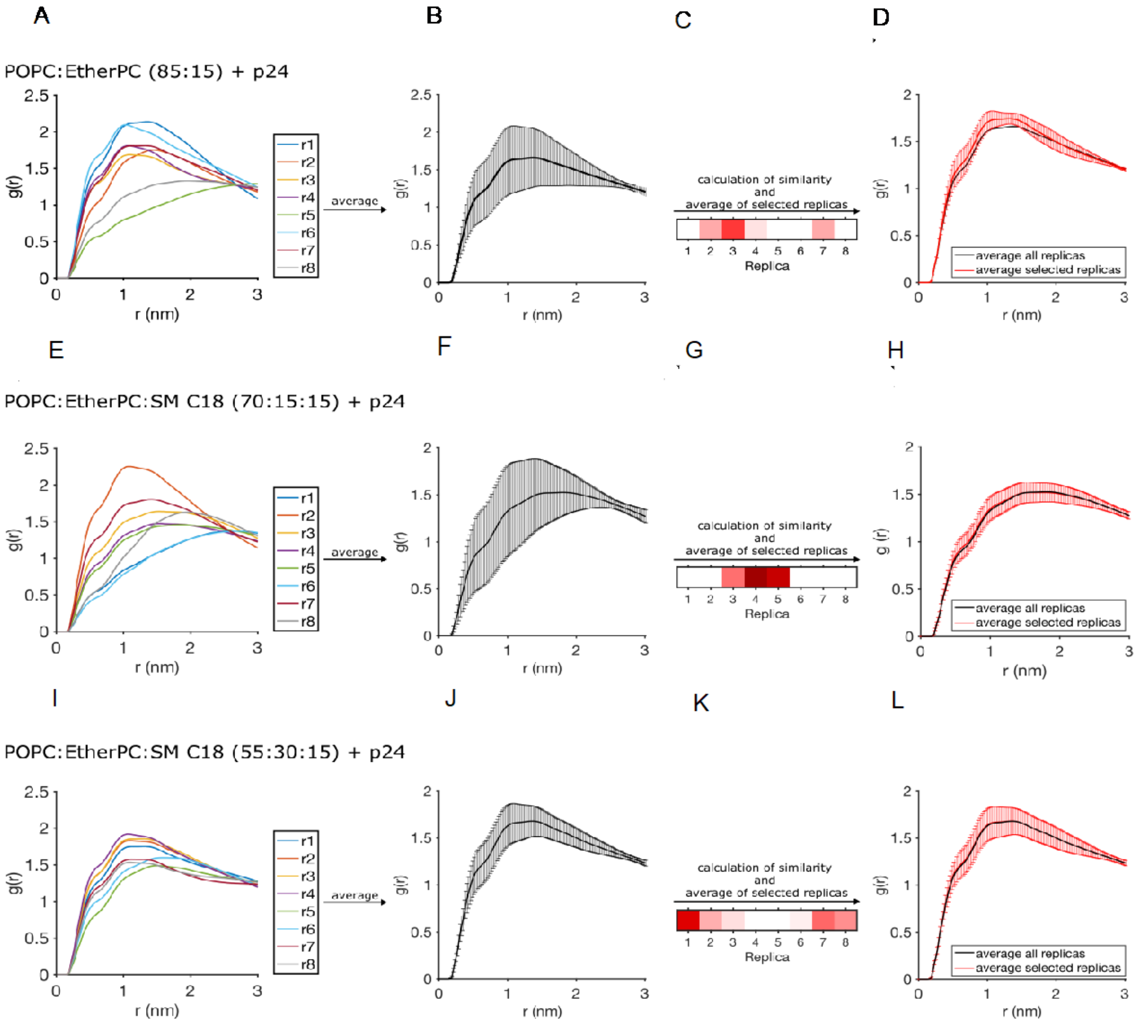


**Supplementary Figure 5. Selection of the converged simulations. RDFs of ether PC with respect to p24 for all the replicas of the system.**

**a,** "POPC:EtherPC 85:15 + p24", **e,** "POPC:EtherPC:SM C18 70:15:15 + p24" and **i**, "POPC:EtherPC:SM C18 55:30:15 + p24". Average curve is obtained by averaging the RDF curves of all the 8 replicas for the systems: **b,** "POPC:EtherPC 85:15 + p24", **f,** "POPC:EtherPC:SM C18 70:15:15 + p24" and **j,** "POPC:EtherPC:SM C18 55:30:15 + p24". Map reporting the similarity values between each replica and the average curve for the systems **c,** "POPC:EtherPC 85:15 + p24", **g,** "POPC:EtherPC:SM C18 70:15:15 + p24" and **k,** "POPC:EtherPC:SM C18 55:30:15 + p24. Dark red indicates very similar curves (Fréchet value close to 0), while white corresponds to non-similarity (Fréchet value above the cutoff of 0.2). Curve obtained averaging the selected replicas (red) and the average of all the replicas for the systems (black) for the systems **d,** "POPC:EtherPC 85:15 + p24", **h,** "POPC:EtherPC:SM C18 70:15:15 + p24" and **l,** "POPC:EtherPC:SM C18 55:30:15 + p24".
